# Asymmetric distribution of centromere proteins between germline stem and daughter cells maintains a balanced niche in Drosophila males

**DOI:** 10.1101/2022.09.29.510087

**Authors:** Antje M Kochendoerfer, Elaine M Dunleavy

## Abstract

In the germline, stem cells undergo asymmetric cell division (ACD) to give rise to one new stem cell as well as one daughter cell that differentiates and eventually gives rise to the gamete (egg or sperm). The silent sister hypothesis proposes the selective inheritance of one set of sister chromatids carrying specific epigenetic marks to explain how genetically identical stem and daughter cells can adopt different fates. It also proposes that centromeres of sister chromatids might differ epigenetically. In Drosophila germline stem cells (GSCs), the centromeric histone CENP-A - CID in flies - is asymmetrically distributed between sister chromatids such that chromosomes that end up in the GSC harbour more CID at centromeres. In this system, a model of ‘mitotic drive’ has been proposed. According to this model, stronger and earlier centromere and kinetochore interactions with microtubules bias sister chromatid segregation during ACD. Here we show that in Drosophila males, CID, CENP-C and CAL1 are asymmetrically distributed between newly divided GSCs and daughter cells that have entered S-phase of the next cell cycle. We find that overexpression of CID, overexpression of CID together with CAL1 or CENP-C depletion disrupts CID asymmetry, with an increased pool of GSCs relative to daughter cells detectable in the niche. This result suggests a shift toward GSC self-renewal rather than differentiation, which is important to maintain tissue homeostasis. Over-expression of CAL1 does not disrupt asymmetry, nor does it affect the GSC and daughter cell balance, but instead drives germ cell proliferation in the niche. Our results in male GSCs are closely aligned with previous observations in female GSCs, indicating that despite differences in signaling, organisation and niche composition, centromere asymmetry and its effects on GSC maintenance are conserved between the sexes.

## Introduction

Asymmetric cell division (ACD) in stem cells generates two daughter cells with unequal fates; one self-renewing daughter cell to replenish the stem cell pool and another daughter cell that differentiates (Sunchu & Cabernard, 2020). ACD is important for maintaining tissue homeostasis and its mis-regulation can lead to several diseases, including cancer and infertility (Knoblich, 2010; Morrison & Kimble, 2006). The silent sister hypothesis (SSH) was put forward as a mechanism to explain how ACD can give rise to genetically identical daughter cells with two different fates. This hypothesis proposed the existence of epigenetically distinct sister chromatids, which could be selectively inherited between stem and daughter cells (Lansdorp, 2007). Furthermore, it proposed that distinct epigenetic marks on sister chromatids can result in differential gene expression in the two daughter cells accounting for different cell fates. Evidence from germline stem cells (GSCs) in Drosophila males have provided an initial support for the SSH (Tran *et al*, 2012; Xie *et al*, 2015). These studies demonstrated a bias in the incorporation and inheritance of pre-existing ‘old’ histones and newly synthesized ‘new’ histones (and associated post-translational modifications) in GSCs undergoing ACD. More recently, differential condensation of sister chromatids via the pre-replication complex component Cdc6 was also reported in GSCs (Ranjan *et al*, 2022). Critically, disruption of histone inheritance patterns leads to stem cell maintenance and cellular differentiation phenotypes in Drosophila testes (Ranjan *et al*., 2022; Tran *et al*., 2012; Xie *et al*., 2015).

An additional element of the SSH proposed the existence of epigenetically distinct sister centromeres (Lansdorp, 2007). Evidence from Drosophila intestinal stem cells provided a first support for this notion (García Del Arco *et al*, 2018). In this study, labelling and tracking of ‘old’ and ‘new’ versions of the centromeric histone CENP-A (called Centromere Identifier CID in flies) showed that ‘old’ CENP-A was preferentially retained in the stem cell. More recent experiments performed in Drosophila GSCs have now shown that in mitosis, centromeres of sister chromatids that end up in the stem cell harbour a higher level of CID compared to those that end up in the differentiating daughter cell (Dattoli *et al*, 2020; Ranjan *et al*, 2019). In male GSCs, an approximate 1.5-fold asymmetric distribution of CID between sister centromeres was measured at prophase/prometaphase, anaphase and telophase (Ranjan *et al*., 2019). In female GSCs, 1.2-fold CID asymmetry was measured at prometaphase/metaphase (Dattoli *et al*., 2020). In addition to CID, Drosophila centromeres are comprised of two major components, the centromere assembly factor Chromosome Alignment 1 (CAL1) (Chen *et al*, 2014) and Centromere Protein CENP-C that serves as the platform for kinetochore assembly and microtubule attachment (Przewloka *et al*, 2011). In male GSCs, an approximate 1.7-fold asymmetry for CAL1 was reported at prometaphase (Ranjan *et al*., 2019), while in females 1.4-fold asymmetry for CENP-C was measured at anaphase and telophase (Carty *et al*, 2021). Also in males, the kinetochore component NDC80 displayed an overall 1.49-fold enrichment in the sister chromatids segregated to the stem cell compared to those segregated to the daughter cell (Ranjan *et al*., 2019). Spindle microtubules are also asymmetrically distributed in GSCs. In males, there is a temporal asymmetry in microtubule dynamics such that centromeres on the ‘stem cell’ side, with more CID, are captured first (Ranjan *et al*., 2019); in females, more spindle microtubules were found to emanate from the pole positioned in the direction of the future stem cell (Dattoli *et al*., 2020). Based on these observations, a mitotic drive model has been proposed as a mechanism that contributes to ACD via centromere strength (Kochendoerfer *et al*, 2021; Ranjan & Chen, 2022). According to this model, during ACD, a cascade of spatial and temporal events occurs in an asymmetric fashion (centrosome and microtubule activity, nuclear membrane breakdown, centromere and kinetochore assembly). Ultimately, these events bias which sister chromatid ends up in the stem or daughter cell that in turn sets up cell fate differences through the inheritance of a distinct set of epigenetic marks.

To determine the importance of centromere asymmetry for ACD in GSCs, and ultimately in cell fate, studies so far have focused on centromere depletion and overexpression strategies. Overexpression of CID together with CAL1 in female GSCs disrupted CID asymmetry leading to more stem cells relative to daughter cells in the ovary; whereas CID or CAL1 knockdown in GSCs blocked cell division (Dattoli *et al*., 2020). While overexpression of CENP-C was not sufficient to disrupt centromere asymmetry in female GSCs, knockdown of CENP-C also resulted in more GSCs relative to daughter cells in the ovary (Carty *et al*., 2021). CAL1 knockdown experiments in male GSCs have shown loss of CID asymmetry and an eventual reduction in GSC number (Ranjan *et al*., 2019). However, the impact of CAL1 depletion on the balance of stem and daughter cells in the male niche was not assessed. Moreover, CENP-C asymmetry and function in male GSCs is so far unexplored. In this study, we investigate effects of CID, CAL1, CENP-C depletion and overexpression on centromere asymmetry in male GSCs and impacts on the balance of stem and daughter cells in the niche.

## Results

### CID, CAL1 and CENP-C are asymmetrically distributed between GSCs and GBs at S-phase

In Drosophila males, the GSC niche is located at the apical end of each testis. Depending on the fly line, between 6-15 GSCs are positioned surrounding a cluster of non-dividing stromal cells called the hub (Kochendoerfer *et al*., 2021). GSCs frequently undergo ACD to give rise to a new GSC that remains associated with the hub, as well as a daughter cell – the gonialblast (GB) – that moves away from the hub and initiates differentiation (Figure 1A). GBs undergo four rounds of mitosis to generate 2-, 4-, 8- and 16-cell cysts (CC) that undergo meiosis to eventually generate 64 haploid spermatozoa. GSCs or GBs can be identified based on the spectrosome, which is a spectrin-rich cytoplasmic organelle. The spectrosome changes morphology through the cell cycle and it can be used to indicate GSC cell cycle phase (Ables & Drummond-Barbosa, 2013). For example, newly divided GSCs have a very short G1-phase and at the end of mitosis both GSCs and GBs enter synchronously into S-phase, connected via a bridge-shaped spectrosome.

**Figure 1.**
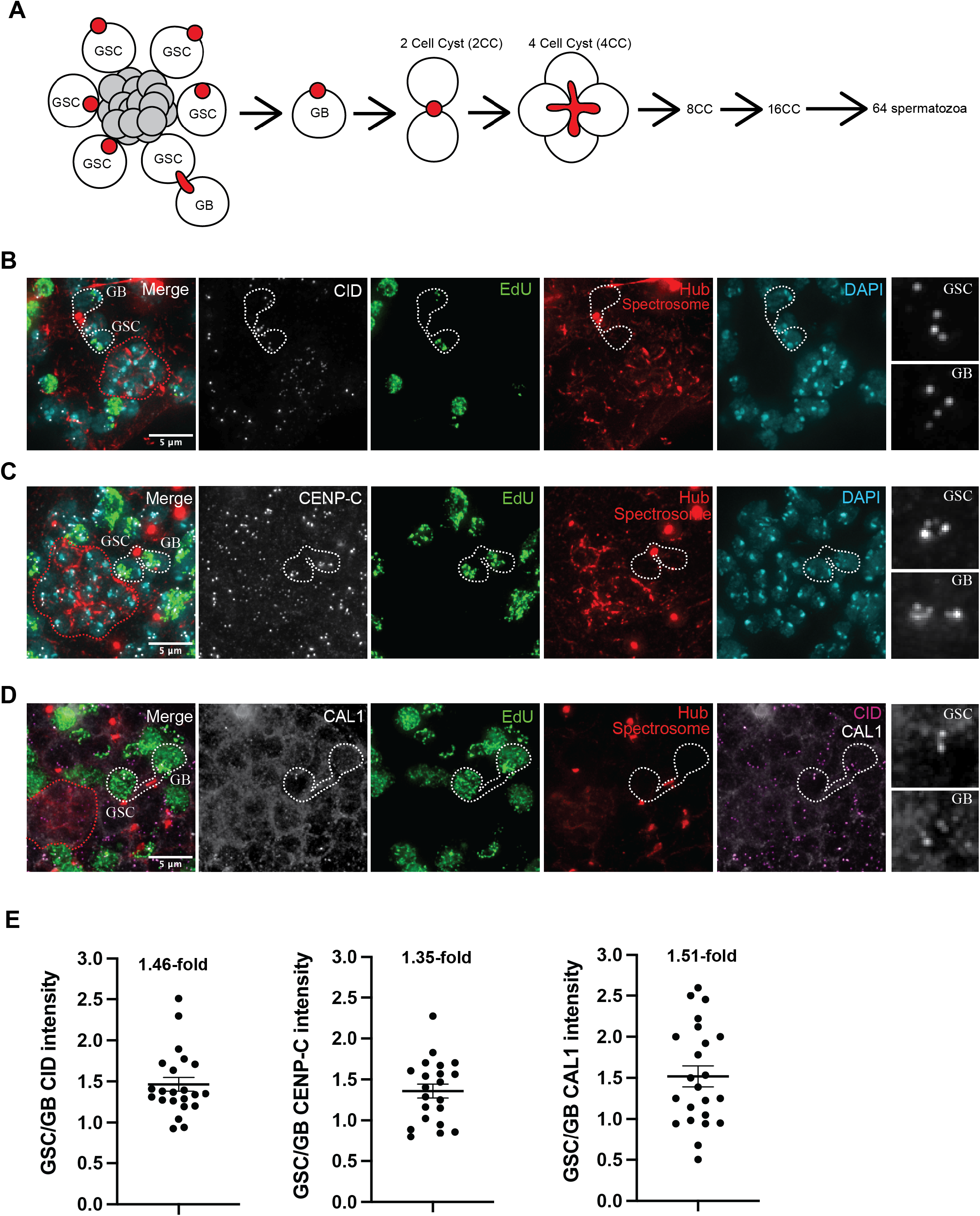
(A) Schematic of the male germline stem cell (GSC) niche and cell divisions occurring in the Drosophila adult testis. GSCs are positioned in close contact with a patch of condensed nuclei that comprise the hub (grey circles). GSCs and daughter gonialblasts (GBs) can be identified by a round spectrin-rich organelle, the spectrosome. The spectrosome forms a bridged shape in S-phase that connects newly divided GSC and GB pairs. GBs undergo four rounds of mitosis generating 2-, 4-, 8-and 16-cell cysts (CC). 16CC undergo meiosis to give bundles of 64 haploid spermatozoa. (B, C) Immunofluorescent image of the GSC niche of wild type Drosophila testes stained for CID (B) or CENP-C (C) (white), the spectrosome together with the hub (red), DAPI (cyan) and pulse labelled with EdU (green). Scale bar = 5 μm. Red dashed line outlines the hub. White dashed line highlights a GSC and GB pair in S-phase; zoom shows CID (B) or CENP-C (C) foci in highlighted GSC and GB nuclei. (D) Immunofluorescent image of the GSC niche of wild type Drosophila testes stained for CAL1 (white), the spectrosome together with the hub (red), CID to mark centromeres (magenta) and pulse labelled with EdU (green). Scale bar = 5 μm. Red dashed line outlines the hub. White dashed line highlights a GSC and GB pair in S-phase; zoom shows CAL1 foci in highlighted GSC and GB nuclei. Scale bar = 5 μm. (E) Quantitation of the ratio of total CID, CENP-C or CAL1 fluorescent intensity (integrated density) between GSC and GB S-phase pairs in the wild type. Each point represents the ratio of total intensity between GSC versus its corresponding GB. Values 1.46 (for CID), 1.35 (for CENP-C), 1.51 (for CAL1) indicate mean fold differences in intensity. At least 20 GSC-GB pairs were analysed per quantitation.

Previous studies showed that in Drosophila male GSCs, CID and CAL1 are asymmetrically distributed between the centromeres of sister chromatids in mitosis (Ranjan *et al*., 2019). To assess whether this asymmetry is maintained in the next cell cycle, we pulse labelled testes with EdU to identify cells undergoing DNA replication in S-phase and used immunolabelling of the bridge-shaped spectrosome to identify newly divided GSC-GB pairs. Immunofluorescent staining for CID, CENP-C and CAL1 indicated that more centromere proteins are present in the GSC compared to the GB (Figure 1B, 1C, 1D). Quantitation showed an approximate 1.46-fold enrichment for CID and a 1.35-fold CENP-C in GSCs compared to GBs (Figure 1E). CAL1 showed 1.51-fold enrichment in GSCs compared to GBs, although we noted that measured intensities were more variable for CAL1, which might reflect its dynamics at this cell cycle time. These observations are in line with measurements of CID level on sister chromatids in male GSCs performed at pro-metaphase/anaphase/telophase in which a 1.5-fold enrichment of CID on the sister chromatids that end up in the GSC compared to those that end up in the GB (Ranjan *et al*., 2019). Our experiments now show that the asymmetric distribution of centromere proteins CID, CENP-C and CAL1 between sister chromatids first observed in mitosis, is carried through into S-phase of the next cell cycle.

### RNAi depletion of CID, CAL1 or CENP-C leads to a reduced number of GSCs per testis

We next aimed to determine if the asymmetric distribution of CID, CENP-C and CAL1 observed at S-phase is important for GSCs maintenance. Previous knockdown experiments in male GSCs showed that CAL1 depletion disrupted the asymmetric CID distribution between sister chromatids at anaphase/early telophase (Ranjan *et al*., 2019). A reduced number of GSCs per testis was also reported upon CAL1 RNAi. Using a similar strategy (spatiotemporally controlled knockdown using nanos-GAL4 activated by the tubGAL80 temperature sensitive driver, *nanos-GAL4; tubGAL80ts*), we performed knockdowns of CID and CENP-C, in addition to CAL1 knockdown with an independent RNAi line. We confirmed respective knockdowns by immunofluorescent staining of testes for CID, CAL1 or CENP-C 10 days after RNAi induction (Supp Figure 1A). We next utilised staining for phosphorylated mothers against Decapentaplegic (pMAD), a Bone Morphogenetic Protein (BMP) signaling indicator present in GSCs (Song *et al*, 2004), together with the presence of the round spectrosome in cells in direct contact with the hub to identify and quantify GSC number per testis. Using these criteria, we counted between 4 and 5 GSCs per testis in *nanos-GAL4; tubGAL80ts* and in non-target mCherry RNAi controls 10 days after RNAi induction (Figure 2A). In the CID, CAL1 and CENP-C RNAi, the number of GSCs per testis was significantly reduced to approximately 2 for each line. This result indicates that depletion of CAL1, CID or CENP-C in GSCs leads to a reduced number of GSCs per testis. Despite this reduction in GSCs observed 10 days after RNAi induction, all lines continued to produce mature sperm indicating that the final stage of spermiogenesis was not yet affected at this RNAi timepoint (Supp Figure 1B).

**Figure 2.**
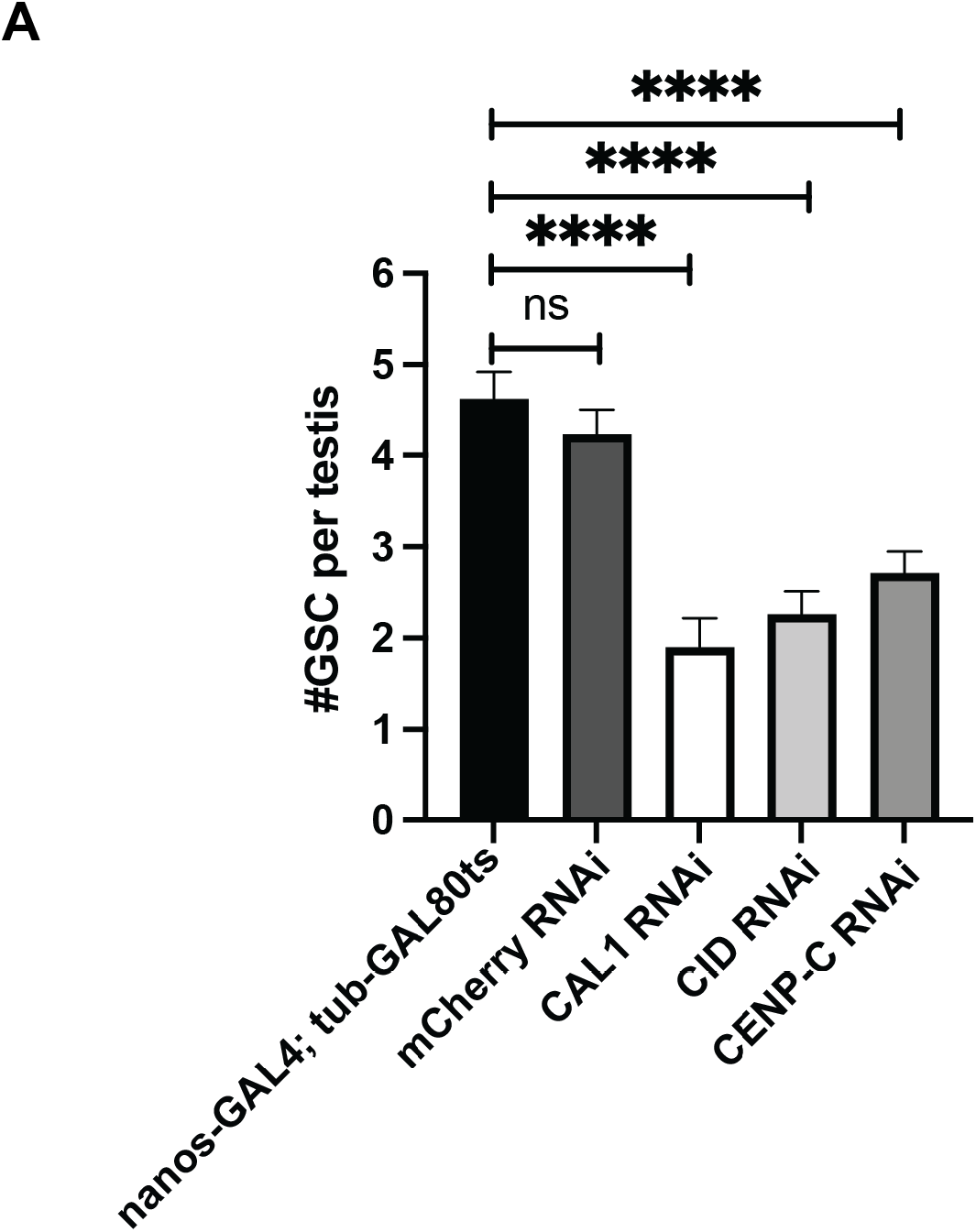
Quantitation of the number of GSCs per testes (n=20-30) in the *nanos-GAL4; tubGAL80ts* control, nontarget mCherry RNAi control, CAL1 RNAi, CID RNAi and CENP-C RNAi 10 days after RNAi induction. GSCs were identified as pMAD-positive cells with a round spectrosome that were attached to the hub. **** p<0.0001, ns non-significant, error bars = SEM.

### Overexpression of CID or CID together with CAL1, but not CAL1 alone, disrupts the asymmetric distribution of CID between GSCs and GBs

The observed GSC loss in testis depleted for CID, CAL1 and CENP-C is indicative of stem cell self-renewal failure. However, this phenotype could also arise due to cell division defects related to these essential genes. As a complementary means to disrupt centromere asymmetry in GSCs, we utilised the GAL4-UAS system to induce the overexpression of centromere components in GSCs. Importantly, we previously employed this strategy to successfully disrupt centromere asymmetry in female GSCs (Dattoli *et al*., 2020). For this purpose, we crossed flies carrying CID-mCherry (CID_OE), CAL1-YFP (CAL1_OE), or both CAL1-YFP and CID-mCherry (CAL1-CID_OE) transgenes to a *nanos-GAL4* driver line. We first validated the induced expression of the transgenes in male GSCs from 3-day old F1 progeny and confirmed expected localisation patterns for CID-mCherry, CAL1-YFP and CID-mCherry_CAL1-YFP (Supp Figure 2A). Next, we stained the overexpression lines with anti-CID antibody to quantify total CID intensity and determine the level of CID overexpression in GSCs (Figure 3A). For this we isolated GSCs with a round spectrosome, which are mostly at G2-phase, the time of CID assembly in these cells (Ranjan *et al*., 2019). Quantitation revealed an approximate 2-fold increase in total CID intensity per GSCs in all three overexpression lines (Figure 3B). Additionally, we quantified the total number of CID foci per GSC nucleus to determine whether CID and/or CAL1 overexpression resulted in extra CID foci per nucleus (Figure 3C). In control GSCs, less than 4 foci were detected indicative of centromere clustering in these cells. In CID_OE and CAL1-CID_OE lines the number of foci is significantly increased to approximately 5, possibly representing ectopic centromeres or reduced centromere clustering, whereas no significant increase was detected in the CAL1_OE. We next assayed for CID asymmetry in GSC-GB pairs in S-phase (Figure 3D). We found that CID or CAL1-CID overexpression disrupts CID asymmetry, giving a symmetric distribution of CID between GSCs and GBs (GCS/GB CID intensity ratio of approximately 1.0) compared to the asymmetric distribution measured in the *nanos-GAL4* control (GCS/GB CID intensity ratio =1.43), which was consistent with the wild type value (Figure 1E). In contrast, CAL1 overexpression did not significantly change CID distribution and the GCS/GB CID intensity ratio remained at 1.48 (Figure 3E). All three overexpression lines produce mature sperm stored in the seminal vesicle indicating that F1 males were likely to be fertile (Supp Figure 2B).

**Figure 3.**
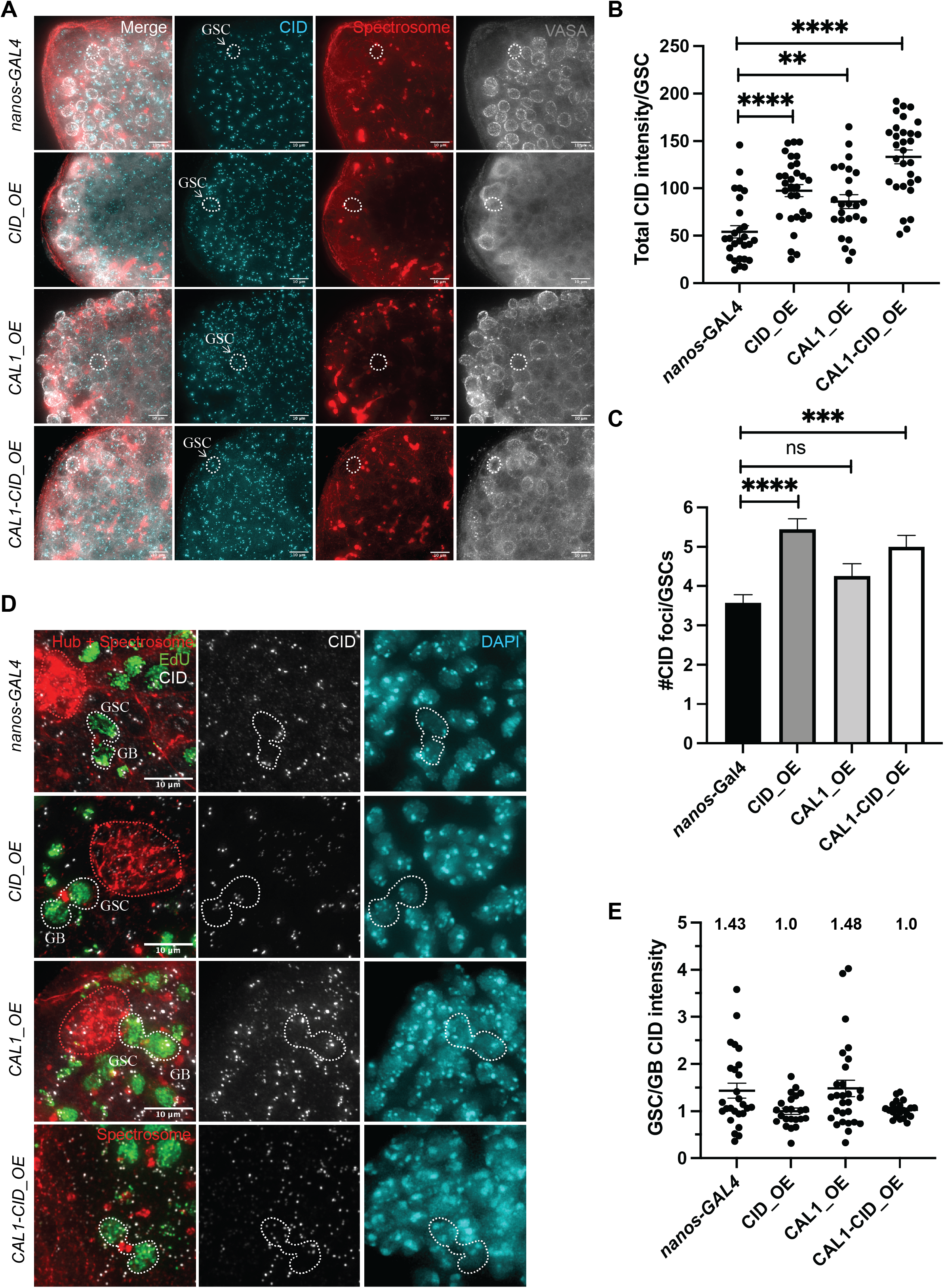
(A) Immunofluorescent image of control *nanos-GAL4 testis*, or testis overexpressing CID-mCherry (CID_OE), CAL1-YFP (CAL1_OE) and CAL1-YFP together with CID-mCherry (CAL1-CID_OE) stained for CID (cyan), the spectrosome (red), and VASA marking germ cells (grey). White arrows and dashed circles indicate typical GSCs isolated for quantitation that were VASA positive, with a round spectrosome and located adjacent to the hub (identified using DAPI, not shown). Scale bar = 10 μm. (B) Quantitation of total CID fluorescent intensity (integrated density) in GSCs (n=25-30) in the control *nanos-GAL4*, and in CID_OE, CAL1_OE or CAL1-CID_OE lines. **** p<0.0001, ** p<0.01, error bars = SEM. n= 30 GSCs per quantitation. (C) Quantitation of total number of CID foci in GSCs (n=28) in the control *nanos-GAL4*, and in CID_OE, CAL1_OE or CAL1-CID_OE lines. **** p<0.0001, *** p<0.001, ns non-significant, error bars = SEM. n= 30 GSCs per quantitation. (D) Immunofluorescent image of control *nanos-GAL4* testis, or testis overexpressing CID-mCherry (CID_OE), CAL1-YFP (CAL1_OE) or CAL1-YFP together with CID-mCherry (CAL1-CID_OE) stained for CID (white), the spectrosome together with the hub (red), DAPI (cyan) and pulse labelled with EdU (green). Red dashed line outlines the hub. White dashed line highlights a GSC and GB pair in S-phase. The cluster of DAPI-dense nuclei was used to identify the hub in CAL1-CID_OE. Scale bar = 10 μm. (E) Quantitation of the ratio of total CID fluorescent intensity (integrated density) between GSC and GB S-phase pairs in *nanos-GAL4*, CID_OE or CAL1-CID_OE lines. Values 1.43 (for *nanos-GAL4*), 1.0 (for CID_OE), 1.0 (for CAL1-CID_OE), 1.48 (for CAL1_OE) indicate mean fold differences in intensity. Between 25-30 GSC-GB pairs were analysed per quantitation, with the exception of CAL1-CID (n=21).

### Overexpression of CID or CID together with CAL1 disrupts the stem and daughter cell balance in the niche; CAL1 overexpression leads to more germ cell proliferation

To determine effects of CID and/or CAL1 overexpression on the balance of stem and daughter cells in the GSC niche, we devised two functional assays (Figures 4 and 5). The first assay was based on spectrosome morphology and VASA staining to mark germ cells (Figure 4A). VASA-positive cells with a round spectrosome located in close association with the hub were annotated as GSCs. In addition, all VASA positive cells with a round spectrosome up to and including the 2-cell cyst stage (2CC) were counted. Compared to the *nanos-GAL4* control, CID_OE, CAL1_OE and CID_CAL1_OE lines showed a disrupted and less organised GSC niche (Figure 4B). An expansion of the hub diameter and number of nuclei was also measured in the case of CID_OE and CAL1_OE (Supp Figure 3A, 3B, 3C), which are additional indicators of a disrupted GSC niche.

**Figure 4.**
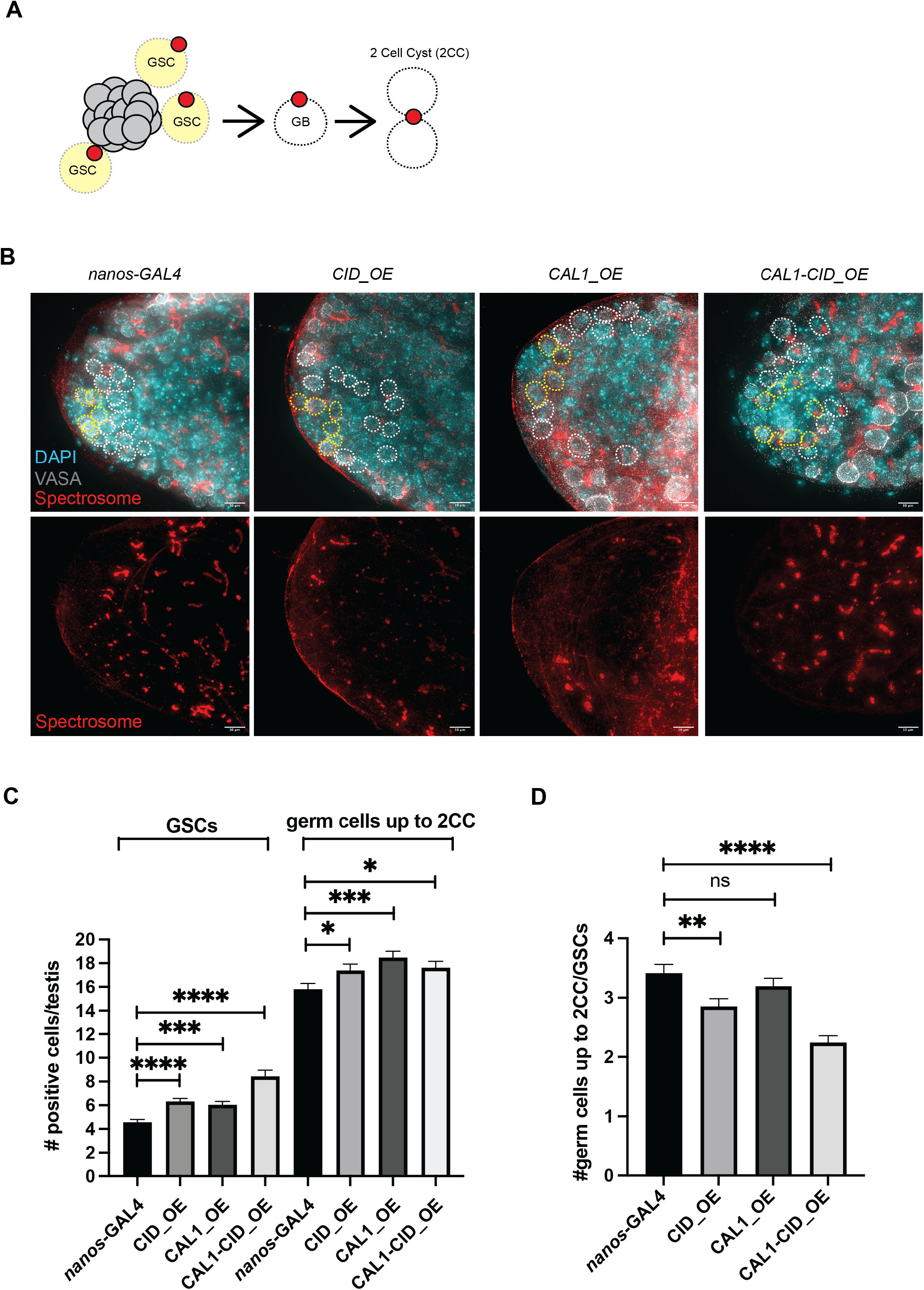
(A) Schematic of the assay used to quantify the balance of GSCs (in yellow), GBs and 2CC (in white) in Drosophila adult testis. (B) Immunofluorescent image of control *nanos-GAL4* testis, or testis overexpressing CID-mCherry (CID_OE) or CAL1-YFP (CAL1_OE) or CAL1-YFP together with CID-mCherry (CAL1-CID_OE) stained for the spectrosome (red), VASA marking germ cells (grey) and DAPI (cyan). Yellow dashed circles indicate GSCs with a round spectrosome located in contact with the hub. White dashed circles indicate GBs or 2CCs with a round spectrosome not in contact with the hub. Scale bar = 10 μm. (C) Quantitation of total number of GSCs or germ cells up 2CC in the control *nanos-GAL4*, and in CID_OE, CAL1_OE or CAL1-CID_OE lines. **** p<0.0001, *** p<0.001, * p<0.05, error bars = SEM. n= 30 testes were analysed per quantitation. (D) Ratio of all germ cells up to 2CC divided by the number of GSCs the control *nanos-GAL4*, and in CID_OE, CAL1_OE or CAL1-CID_OE lines. **p<0.01, **** p<0.0001, ns non-significant, error bars = SEM.

**Figure 5.**
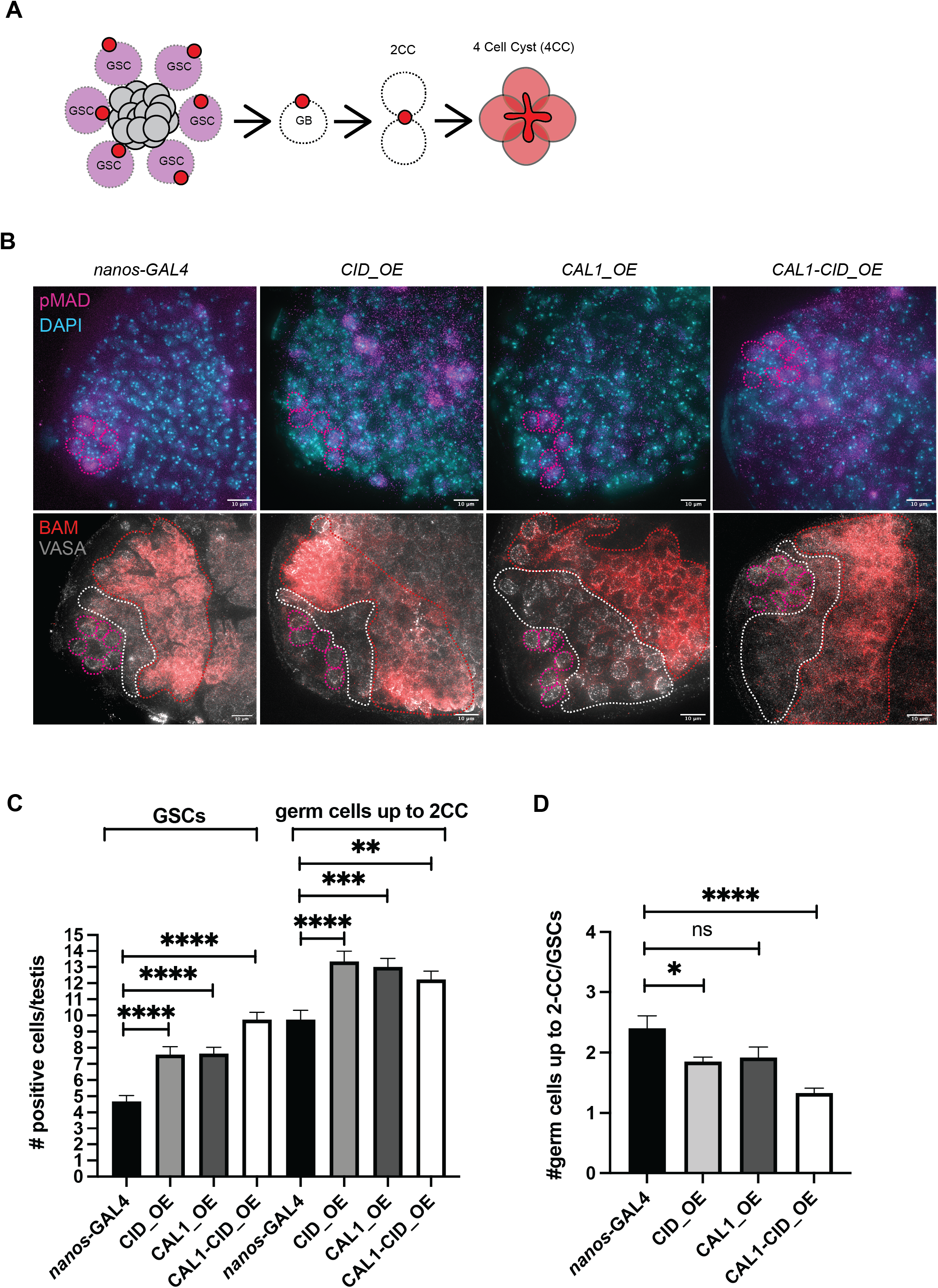
(A) Schematic of the assay used to quantify the balance of GSCs (in magenta), GBs and 2CC (in white) in Drosophila adult testis. GSCs were counted based on a pMAD-positive signal with a round spectrosome adjacent to the hub. All germ cells up to the 2CC were VASA-positive and BAM-negative. BAM expression in 4CC is shown in red. (B) Immunofluorescent image of control *nanos-GAL4* testis, or testis overexpressing CID-mCherry (CID_OE) or CAL1-YFP (CAL1_OE) or CAL1-YFP together with CID-mCherry (CAL1-CID_OE) stained for pMAD (magenta), BAM (red), VASA marking germ cells (grey) and DAPI (cyan). Magenta dashed circle indicate pMAD positive GSCs in contact with the hub. White dashed circles indicate VASA positive cells counted up to 2CC. Scale bar = 10 μm. (C) Quantitation of total number of GSCs or germ cells up the 2CC in the control *nanos-GAL4*, and in CID_OE, CAL1_OE or CAL1-CID_OE lines. **** p<0.0001, *** p<0.001, **p<0.01, ns non-significant, error bars = SEM. n= 30 testes were analysed per quantitation. (D) Ratio of all germ cells up to 2CC divided by the number of GSCs the control *nanos-GAL4*, and in CID_OE, CAL1_OE or CAL1-CID_OE lines. **** p<0.0001, * p<0.05, ns non-significant, error bars = SEM.

Using our criteria to identify GSCs, we quantified the number of GSCs per testis (Figure 4C). Between 4 and 5 GSCs were detected in the control, whereas this number increased in CID_OE (6.3), CAL1_OE (6) and CAL1_CID_OE (8.4) lines. Quantitation of all VASA positive cells with a round spectrosome up to 2CC also revealed an increase in CID_OE (17.3), CAL1_OE (18.4) and CID_CAL1_OE (17.6) lines compared to the control (15.8). When expressed as a ratio, for every 1 GSC we observed approximately 3.4 ‘daughter cells’ in the control (Figure 4D). In CID_OE and CAL1-CID_OE, for every 1 GSC there are 2.8 and 2.2 daughter cells respectively, meaning that there are less daughter cells for every stem cell. For CAL1 overexpression, the ratio (3.1) is not significantly different from control, however we observed an overall increase in the total number of germ cells in the niche. Longer term CAL1 overexpression studies (after 10-days) showed a further increase in the total number of round-spectrosome and VASA positive germ cells per testis (Supp Figure 3D). In this case, for every 1 GSC, 4.5 daughter cells were detected (Supp Figure 3E). Taken together, results for CAL1 overexpression after 3 and 10 days indicate that CAL1 expression might drive germ cell proliferation, including GSCs, in the niche.

To validate these findings, we developed a second functional assay to measure the balance of stem and daughter cells in the male GSC niche based on an independent set of cellular markers (Figure 5A). In these experiments, GSCs were identified based on pMAD signal and contact with the DAPI-dense nuclei that comprise the hub region. In addition, all VASA positive cells that were negative for the differentiation marker *bag of marbles* (BAM) that is turned on at the 4-cell cyst (4CC) stage. Again, we noted that compared to the *nanos-GAL4* control, CID_OE, CAL1_OE and CAL1-CID_OE lines showed a disrupted and less organised GSC niche (Figure 5B). Ectopic pMAD signal was also noted, particularly in CID_OE and CAL1-CID_OE lines, which further indicates disrupted cell signaling in these niches. Using this second set of criteria to identify GSCs, we quantified the number of GSCs per testis (Figure 5C). Approximately 5 (4.6) GSCs we detected in the control, whereas this number increased in CID_OE (7.5), CAL1_OE (7.6) and CAL1-CID_OE (9.7) lines. Quantitation of all VASA positive cells up to the 4CC when BAM is expressed also revealed an increase in CID_OE (13.3), CAL1_OE (13) and CAL1-CID_OE (12.2) lines compared to the control (9.7). When expressed as a ratio, we observed approximately 2.5 daughter cells for every GSC in the control (Figure 4D). In CID_OE and CAL1-CID_OE, for every 1 GSC there are approximately 1.5 daughter cells, meaning that there are less daughter cells for every stem cell. For CAL1 overexpression, the ratio is not significantly different from the control, however we observe an overall increase in the total number of germ cells in the niche and more variability in these samples (see SEM error bars).

Taken together, both assays of stem and daughter cell balance indicate that upon CID_OE or CAL1-CID_OE the number of daughter cells relative to GSCs is reduced meaning that the pool of GSCs is increased. In the case of CAL1_OE, both assays indicate no change in the balance of stem to daughter cells. Instead, the overall total number of germ cells (GSCs up to 2CCs) is increased upon CAL1 overexpression.

### Extra GSCs observed upon CID/CAL1 overexpression are not due to dedifferentiation

Given that we detected an increased pool of GSCs upon CID and CAL1-CID overexpression, we next wanted to determine the origin of such cells. One possibility is that extra GSC-like cells have arisen due to dedifferentiation of GBs, as previously documented (Cheng *et al*, 2008). Centrosome positioning can be used to assay for GSC-like cells derived from dedifferentiated GBs (Cheng *et al*., 2008). We stained CID_OE, CAL1_OE and CAL1-CID_OE lines with the centrosome marker anti-gamma tubulin and calculated the percentage of GSCs with misorientated centrosomes per testis. GSCs in which one centrosome is normally positioned adjacent to the hub interface are defined as oriented; whereas GSCs with both centrosomes positioned away from the hub are defined as misoriented (Supp Figure 3F). Quantitation in CID_OE (8%), CAL1_OE (3%) and CAL1-CID_OE (10%) lines did not reveal any significant increase in the percentage of GSCs with misoriented centrosomes compared to the *nanos-GAL4* control (6%), suggesting that extra GSCs are not derived from GB dedifferentiation (Supp Figure 3G).

### Overexpression of CENP-C does not affect niche composition

Previous experiments in Drosophila female GSCs showed that in contrast to CID and CAL1 overexpression, overexpression of CENP-C did not impact on centromere asymmetry or cell fate (Carty *et al*., 2021). To investigate whether this observation holds true in the differentially regulated GSC niche in males, we overexpressed HA-tagged CENP-C (HA-CENP-C_OE) in GSCs in testes using the *nanos-GAL4* driver (Figures 6 and 7). We first confirmed the induction of HA-CENP-C in pMAD-positive GSCs using an anti-HA antibody (Supp Figure 4A). We also confirmed that HA-CENP-C localised at centromeres by co-staining with anti-CID antibody (Supp Figure 4B). For GSCs captured at S-phase, no significant change in CID intensity was detected upon HA-CENP-C expression compared to the *nanos-GAL4* control (Figure 6A, 6B), indicating that excess CENP-C is not sufficient to drive centromere assembly. Next, we assessed effects of HA-CENP-C overexpression on CID asymmetry (Figure 6C). Again, we measured no significant change (ratio GSC/GB = 1.43) compared to the *nanos-GAL4* control (ratio GSC/GB = 1.43) (Figure 6C). Finally, we quantified GSCs (pMAD-positive with a round spectrosome and in contact with the hub) and germ cells up to 2CC (VASA-positive with a round spectrosome) (Figure 7). For HA-CENP-C_OE, we did not find a significant change in stem and daughter cell balance compared to the *nanos-GAL4* control (Figure 7B, 7C). This result is very comparable to that reported in female GSCs, in which the overexpression of CENP-C did not affect CID assembly, asymmetry or stem and daughter cell balance in the ovary (Carty *et al*., 2021).

**Figure 6.**
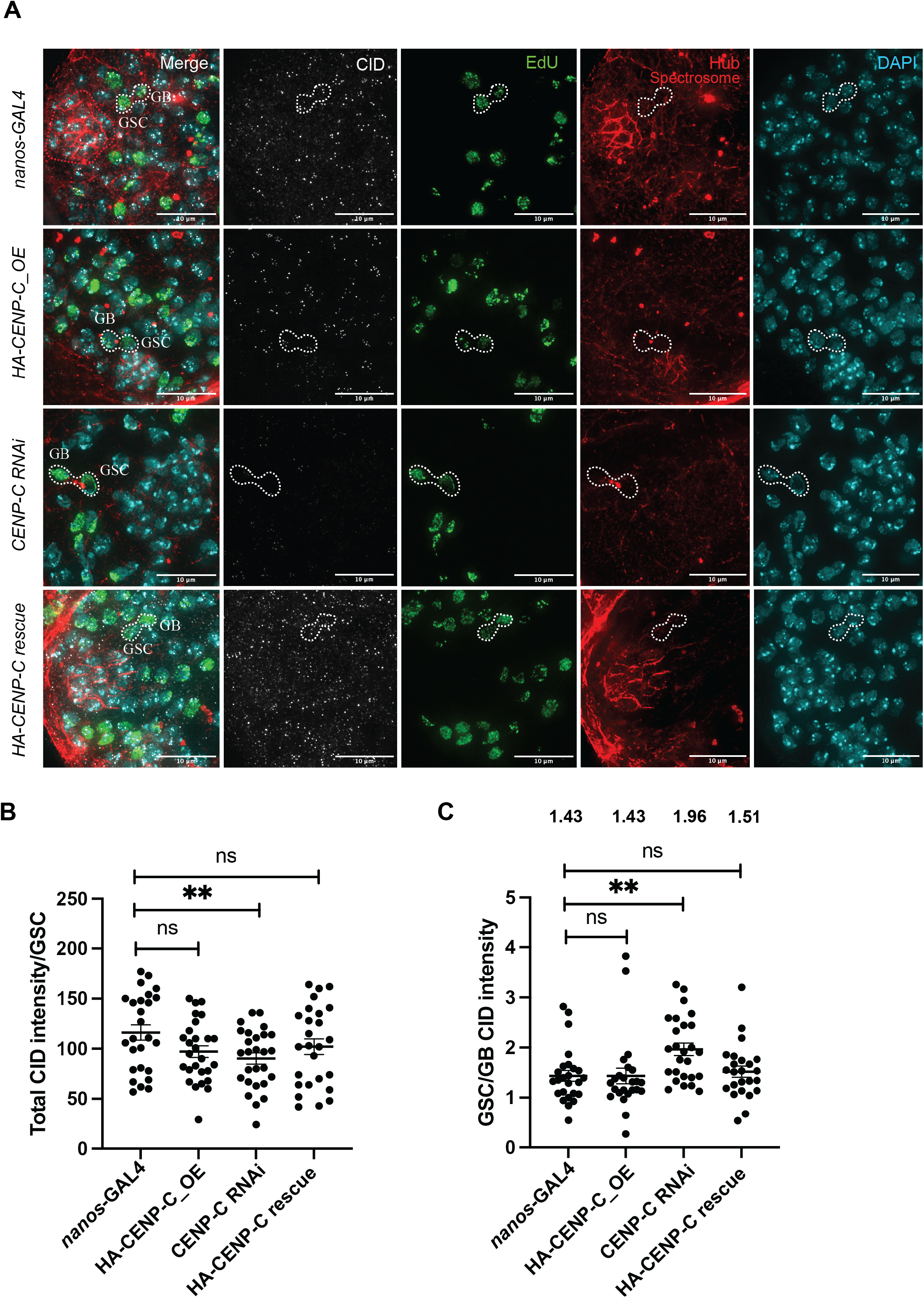
(A) Immunofluorescent image of control *nanos-GAL4*, HA-CENP-C_OE, CENP-C RNAi and HA-CENP-C; CENP-C RNAi (HA-CENP-C rescue) testis stained for CID (white), the spectrosome together with the hub (red), DAPI (cyan) and pulse labelled with EdU (green). White dashed line highlights a GSC and GB pair in S-phase. Scale bar = 10 μm. (B) Quantitation of total CID fluorescent intensity (integrated density) in GSCs in the control *nanos-GAL4*, HA-CENP-C_OE, CENP-C RNAi and HA-CENP-C; CENP-C RNAi (HA-CENP-C rescue) lines. ** p<0.01, ns non-significant, error bars = SEM. n= 25-30 GSCs per quantitation. (C) Quantitation of the ratio of total CID fluorescent intensity (integrated density) between GSC and GB S-phase pairs in *nanos-GAL4*, HA-CENP-C_OE, CENP-C RNAi and HA-CENP-C; CENP-C RNAi (HA-CENP-C rescue) lines. Values 1.43 (for *nanos-GAL4*), 1.43 (for CENP-C_OE), 1.96 (for CENP-C RNAi) and 1.51 (for HA-CENP-C rescue) indicate mean fold differences in intensity. **p<0.01, ns non-significant, error bars = SEM. n= 25-30 GSCs per quantitation.

**Figure 7.**
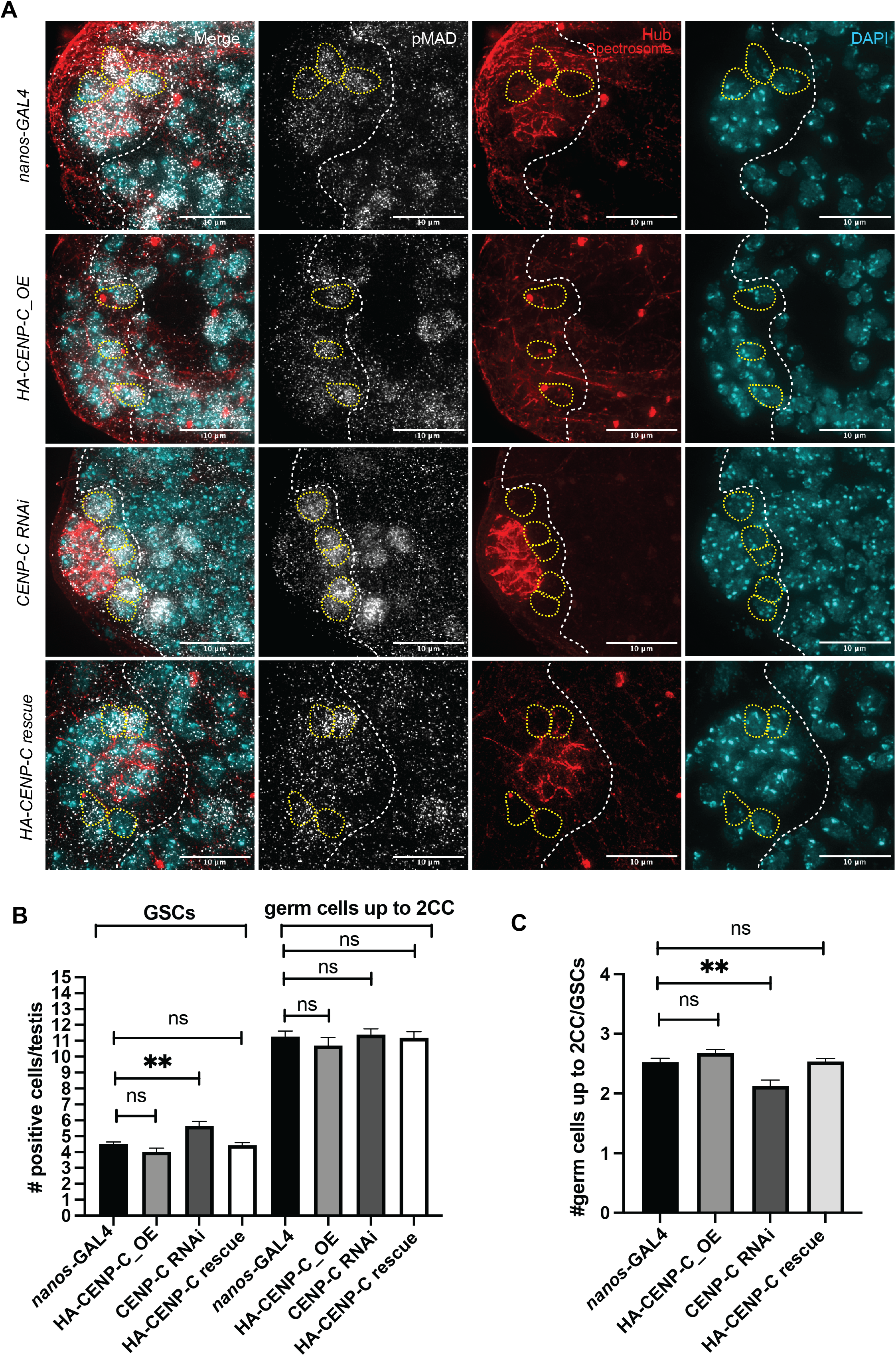
(A) Immunofluorescent image of control *nanos-GAL4*, HA-CENP-C_OE, CENP-C RNAi and HA-CENP-C; CENP-C RNAi (HA-CENP-C rescue) testis stained for pMAD (grey), the spectrosome together with the hub (red) and DAPI (cyan). Yellow dashed circles indicate GSCs with a round spectrosome located in contact with the hub. White dashed line indicates area in which cells with a round spectrosome up to 2CC were counted. Scale bar = 10 μm. (B) Quantitation of total number of GSCs or germ cells up the 2CC in the control *nanos-GAL4*, HA-CENP-C_OE, CENP-C RNAi and HA-CENP-C; CENP-C RNAi (HA-CENP-C rescue) lines. **p<0.01, ns non-significant, error bars = SEM. n= 30 testes were analysed per quantitation. (C) Ratio of all germ cells up to 2CC divided by the number of GSCs in the control *nanos-GAL4*, HA-CENP-C_OE, CENP-C RNAi and HA-CENP-C; CENP-C RNAi (HA-CENP-C rescue) lines. **p<0.01, ns non-significant, error bars = SEM.

### CENP-C depletion disrupts the balance of stem and daughter cells in the niche

While RNAi knockdown of CENP-C using the *nanos-GAL4; tubGAL80ts* driver impacted on the total number of GSCs per testis (Figure 2), we were unable to detect a reduction in CID intensity (Supp Figure 1A). We reasoned that in this RNAi experiment CENP-C kinetochore function might be perturbed, affecting cell division without impacting on CENP-C’s function in CID assembly or maintenance. Instead, CENP-C knockdown driven by the *nanos-GAL4* driver, which is active earlier in the development of the F1 progeny, led to a significant reduction in CID intensity at GSC centromeres (Figure 6A, 6B, Supp Figure 4B). In this CENP-C RNAi experiment, rather than a loss of CID asymmetry, we observed an unexpected increase in CID asymmetry in S-phase GSC-GB pairs (ratio GSC/GB = 1.96) (Figure 6C). Furthermore, the number of GSCs increased (pMAD-positive with round spectrosome and in contact with the hub) with no change in the number of germ cells up to 2CC (VASA positive with round spectrosome) (Figure 7A, 7B, 7C). This resulted in a significant change in the stem and daughter cell balance, with approximately 2 daughter cells for every GSC, compared to 2.5 daughter cells for every GSC in the *nanos-GAL4* control (Figure 7B, 7C). Using the *nanos-GAL4* driver, neither CID asymmetry nor stem and daughter cell balance was affected in the non-target mCherry RNAi control (Supp Figure 4C, 4D). Rescue experiments in which we overexpressed HA-CENP-C that is resistant to the shRNA in the CENP-C RNAi background showed no significant changes in CID intensity at centromeres, CID asymmetry (1.51) nor in the stem to daughter cell balance (2.5:1) (Figures 6 and 7). Finally, CENP-C over-expression or depletion did not appear to impact on fertility, with mature sperm detected in testes from these lines (Supp Figure 4E). These findings for CENP-C in male GSCs are strikingly similar to those reported in female GSCs, in which CENP-C RNAi increases CID asymmetry between GSCs and cystoblasts, the GB equivalent in females. Moreover, CENP-C RNAi in female GSCs shifted the balance of stem and daughter cells such that more GSCs were detectable in the ovary (Carty *et al*., 2021).

## Discussion

This study investigates CID, CAL1 and CENP-C inheritance in newly divided stem (GCS) and daughter (GB) cells that have entered S-phase of the next cell cycle. We find an approximate 1.4-fold asymmetric distribution for CID, CAL1 and CENP-C, with more centromere proteins detectable in GSCs compared to GBs. This result is consistent with previous measurements of CID and CAL1 asymmetry at mitosis in male GSCs (Ranjan *et al*., 2019), as well as asymmetries reported for CID and CENP-C at S-phase in female GSCs (Carty *et al*., 2021; Dattoli *et al*., 2020). It is now important to understand when and how such asymmetries are established and maintained during the cell cycle. One possibility is that asymmetry is established during the time of centromere assembly, occurring at G2-phase up to prophase in GSCs. Differential pools of old and new CAL1 at sister centromeres in GSCs supports asymmetric CID assembly and maintenance at this cell cycle time (Ranjan *et al*., 2019). Asymmetric CID assembly in G2-phase might also explain the unique cell cycle timing of centromere assembly in GSCs (Dattoli *et al*., 2020). In these stem cells undergoing ACD, assembly occurs prior to chromosome segregation, as opposed to after in most other (symmetrically diving) cell types (Carty & Dunleavy, 2020). A second - and not mutually exclusive-possibility is that the redistribution of parental CID nucleosomes during S-phase sets up this asymmetry between sister centromeres. Indeed, unidirectional DNA replication forks appear to be enriched in GSCs which might provide a mechanism for this (Wooten *et al*, 2019). Further single-cell investigations in S-phase GSCs in which sister centromeres might be resolved are required to determine if this is the case.

To disrupt centromere asymmetry in GSCs, we employed overexpression and depletion of CID, CAL1 and CENP-C. Spatiotemporally controlled knockdown of CID, CAL1 and CENP-C resulted in reduced numbers of GSCs per testis 5-days after RNAi induction, which is indicative of stem cell self-renewal failure or defective GSC maintenance. Our result for CAL1 depletion is consistent with GSC loss and decreased expression of the GSC marker Stat92E reported to occur 10-days after RNAi (Ranjan *et al*., 2019). In CENP-C RNAi experiments using the *nanos-GAL4; tubGAL80ts* driver, although we recorded a reduction in GSCs per testis, CID level in GSCs appeared to be unaffected. We reasoned that in this RNAi experiment CENP-C kinetochore function might be perturbed, affecting cell division without impacting on CENP-C’s function in CID assembly or maintenance. Therefore, we utilized the *nanos-GAL4* driver to induce a stronger depletion of CENP-C, which resulted in an approximate 22% drop in CID at GSC centromeres. In this case, we noted an expansion of the GSCs pool. We suppose that CENP-C depletion must reach a threshold before it impacts on CID level at centromeres, which in turn is required to impact on GSC maintenance. Our result in males is in line with previous CENP-C RNAi experiments performed in female GSCs. GSCs in the ovary with a 60% reduction in CENP-C displayed a 35% drop in CID at centromeres with an initial expansion of GSC pool and a subsequent depletion of GSC pool upon long term CENP-C RNAi (Carty *et al*., 2021).

Given that CID, CAL1 and CENP-C are essential genes, we adopted an overexpression strategy to overcome analysis of complex cell division phenotypes related to centromere function. Overexpression of CID, CAL1 and CID together with CAL1 resulted in 2-fold increase in CID intensity in GSCs. For overexpression of CID or CID together with CAL1, but not for CAL1 overexpression alone, this perturbed CID asymmetry and gave a symmetric distribution between stem and daughter cells. To determine effects on stem cell fate, we developed two functional assays using independent sets of cellular markers to measure the effect on stem and daughter cell balance in the male niche. Both assays showed that overexpression of CID or CID together with CAL1 resulted in more stem cells relative to daughter cells. We interpret this result as more stem cell self-renewal occurring in the niche, which we can correlate with a symmetric CID distribution between GSCs and GBs. It is now important to understand the origin of these extra GSC-like cells in the niche that according to our analyses do not appear to have come from de-differentiation of GBs. One possibility is that they are arisen due to extra symmetric cell divisions of GSCs, which could be determined using live imaging of the testis stem cell niche (Greenspan & Matunis, 2017). Overexpression of CAL1 alone did not change the asymmetric distribution of CID between GSCs and GBs nor did it change the GSC and daughter cell balance. Instead, both assays showed that CAL1 overexpression correlated with a general an increase in germ cell number in the niche. It is possible that CAL1 might drive cell proliferation in this context and this function could be independent of its role at centromeres. Indeed, a centromere independent function for CAL1 in terminally differentiated enteroblasts in the Drosophila midgut has already been proposed (García Del Arco *et al*., 2018). In the case of CID overexpression or overexpression of CID together with CAL1, we observed approximately one extra CID focus per GSC that might function as ectopic centromeres. Whether ectopic CID could drive changes in centromere asymmetry, asymmetric chromosome segregation or change gene expression in GSCs remains to be determined. Interesting, no extra CID foci were observed upon CAL1 overexpression, which did not affect asymmetry or the GSC and daughter cell balance. These results in male GSCs are very comparable to our previous findings in female GSCs in which CID or CAL1-CID overexpression resulted in more stem cell self-renewal, while CAL1 overexpression increased germ cell proliferation (Dattoli *et al*., 2020).

Different to CID and CAL1, overexpression of CENP-C did not impact on CID intensity at centromeres and had no effect on CID asymmetry or stem cell maintenance. Instead, knockdown of CENP-C (using the *nanos-GAL4 driver*) increased CID asymmetry between GSCs and GBs. Again, this result in males is strikingly similar to our findings in female GSCs in which CENP-C depletion increased the level of CID asymmetry (Carty *et al*., 2021). Results for CENP-C RNAi indicate that more CID asymmetry can also increase the GSC pool in testes and ovaries. Although centromere depletion and overexpression approaches perturbed the number and balance of germ cells in the niche, adult males continued to produce mature sperm. This observation questions the functional impact of having one less stem cell in the niche. Perhaps it is only under conditions of stress or advanced aging that this deficiency might come into play.

Our results provide additional support for the SSH, as well as mitotic and centromere drive in stem cell ACD. We propose that CID level differentially marks sister chromatids, which leads to non-random sister chromatid segregation. This can ultimately impact on the stem and daughter cell balance and cell fate. This effect is likely to be mediated through changes in gene expression due to extrinsic or extrinsic cell signaling. An alternative idea is that CID incorporation into chromatin might directly change gene expression, which has not yet been explored. Our results also identify conserved features between Drosophila male and female GSCs. The male GSC niche is more complex than in females and is distinct in terms of JAK-STAT/Upd signaling, spectrosome positioning, *Bam* expression pattern and germline-soma interactions (Vidaurre & Chen, 2021). Moreover, although in males inheritance of the mother centrosome is critical for GSC maintenance, in females, GSCs inherit the daughter centrosome (Chen & Yamashita, 2021). Yet, despite these well characterised differences between the sexes, centromere asymmetry and effects on cell fate are conserved between male and female GSCs. Although centromere strength alone is unlikely to be the major determinant of stem cell identity, our results place the centromere as an important player in stem cell ACD.

## Materials and Methods

### Fly stocks and husbandry

Stocks were cultured on standard cornmeal medium (NUTRI-fly) preserved with 0.5% propionic acid and 0.1% Tegosept at 20°C under a 12-hour light-dark cycle. All fly stocks used were obtained from Bloomington Drosophila Stock Centre or Vienna Drosophila Stock Centre (#) unless otherwise stated. The following fly stocks were used: wild type *Oregon-R* (BDSC #25211), *UAS-dcr2; nanos-GAL4* (BDSC #25751), *nanos-GAL4; tub-GAL80ts* (kind gift from Yukiko Yamashita), UAS-CID RNAi (VDRC #102090), UAS-CAL1 RNAi (VDRC #45248), UAS-CENP-C RNAi (BDSC #38917), UASp-HA-CENP-C; *SM6 Cy* (kind gift from Kim S. McKim), HA-CENP-C; UAS-CENP-C-RNAi (Carty et al. 2021), mCherry-RNAi (gift from Gary Karpen). Fly lines used for overexpression of CID, CAL1 and CAL1-CID under UAS control were generated in (Dattoli *et al*., 2020). CENP-C knockdown (and rescue) using the *nanos-GAL4* driver was performed at 22 °C. For knockdown using the *nanos-GAL4; tub-GAL80ts* crosses were set at 18°C and progeny were shifted to 29 °C upon eclosion. HA-CENP-C was induced using *nanos-GAL4* at either 25 °C or at 22 °C for rescue experiments. F_1_ progeny were dissected 3 days after eclosion unless stated otherwise. Results obtained from each experiment rely on three biological replicates, unless otherwise specified.

### Immunofluorescence (IF)

Testes were squashed and snap frozen in liquid Nitrogen. After fixation in 4% PFA for 10 minutes (5 minutes for anti-SMAD3/5), samples were immediately washed in 1XPBS-0.4% Triton-X100 (0.4% PBST). Samples were then blocked in 0.4% PBST with 1% BSA for 2-4 hours at room temperature, incubated with primary antibodies (in blocking buffer) overnight at 4 ^°^C. Samples were then washed in 0.4% PBST for 3x 30 minutes. Secondary antibodies are added (1:500 in blocking buffer) for 2 hours at room temperature in the dark. Samples are again washed 3x 30 minutes in 0.4% PBST followed by addition of DAPI (1:1000) for 15 minutes in 1XPBS.

### EdU Incorporation

Testes were dissected and incubated for 30 min with EdU (0.01 mM, Berry & Associates) in 1XPBS and then fixed as described. After washing in 0.4% PBST, testes were incubated for 30 minutes in the dark with 2 mM CuSO_4_, 300 μM fluorescent azide and 10 mM ascorbic acid.

Samples were then washed with 0.4% PBST for 10 minutes and then blocked and stained as described above.

### Antibodies

For immunostaining, the following primary antibodies were used: rabbit anti-CENP-A (CID) antibody (Active Motif 39719; 1:500), sheep anti-CENP-C (Dattoli *et al*., 2020); 1:2000), rabbit anti-VASA (Santa Cruz sc-30210; 1:500), rat anti-VASA (Developmental Studies Hybridoma Bank (DSHB); 1:500), sheep anti-CAL1 (this study, 1:1000), rabbit anti-SMAD3/5 (pMad) (Abcam ab52903; 1:500), DAPI (1:1000), mouse anti-Armadillo (DSHB N2 7A1, 1:700), mouse anti-1B1 (Hts) DSHB, 1:100), mouse anti-BAM (DSHB, 1:50), mouse anti-FIBRILLARIN (AbCam ab5821, 1:500), mouse anti-HA (Thermofisher 26183, 1:500), mouse anti-gamma-tubulin (AbCam ab44928, 1:500), mouse anti-FASCILLIN III (DSHB 7G10, 1:200), rat anti-mCherry (Chromotek 5F8, 1:200). Sheep anti-CAL1 antibody was raised against CAL1 amino acids 748-979 with a C-terminal His tag produced according to (Bade *et al*, 2014). Species specific Alexa-Fluor 405, 488, 546, 647 secondary antibodies were used at a 1:500 dilution.

### Widefield microscopy

Images of immunostained testes mounted in SlowFade Gold antifade reagent (Invitrogen S36936) were acquired using a DeltaVision Elite microscope system (Applied Precision) equipped with a 100x, 60x or 40x oil immersion UPlanS-Apo objective (NA 1.4). Images were acquired as z-stacks with a step size of 0.2 μm. Fluorescence passed through a 435/48 nm; 525/48 nm; 597/45 nm; 632/34 nm band-pass filter for detection of respectively DAPI, Alexa Fluor 488, mCherry and Alexa Fluor 647 in sequential mode.

### Quantification

For each quantification one cell/testis was considered. Images from a single cell (nucleus) were projected (max intensity) to capture all the centromeres present in the cell at a specific cell cycle phase. Image J software was used to measure fluorescent intensity of CID in the following way: The background was subtracted from the projected image. Threshold was adjusted and the image. Size was adjusted in order to eliminate unwanted objects. Following, the command “analyse particles” was used to select centromeres. Finally, integrated density (MGV*area) from each centromere foci were extracted and used as fluorescent intensity to measure the total amount of fluorescence per nucleus.

### Statistical Analyses

Data distribution was assumed to be normal, but this was not formally tested. P value in each graph shown was calculated with unpaired t test with Welch’s correction. All statistical analysis was performed using Prism 9 software.

## Acknowledgements

Stocks were obtained from the Bloomington Drosophila Stock Center (NIH P40OD018537) or the Vienna Drosophila Resource Centre (VDRC, www.vdrc.at). Antibodies obtained from the Developmental Studies Hybridoma Bank, created by the NICHD of the NIH are maintained at The University of Iowa, Department of Biology, Iowa City, IA 52242.

