## Supplementary Data for "Asymmetric distribution of centromere proteins between germline stem and daughter cells maintains a balanced niche in Drosophila males"

**Supplementary Figure Legends: Kochendoerfer and Dunleavy**

**Supplementary Figure 1**

(A) Immunofluorescent image of testis from *nanos-GAL4; tubGAL80ts* control, nontarget mCherry RNAi control, CID RNAi, CAL1 RNAi and CENP-C RNAi 10 days after RNAi induction. Testes are stained for CID (yellow), the spectrosome together with the hub (red), CAL1 or CENP-C (grey) and DAPI (cyan). For mCherry RNAi, mCherry signal in the red channel was captured (shown in grey). Scale bar = 10 μm.
(B) Fluorescent image of mature sperm (stained for DAPI, cyan) in testes of *nanos-GAL4; tubGAL80ts* control, nontarget mCherry RNAi control, CID RNAi, CAL1 RNAi and CENP-C RNAi 10 days after RNAi induction. Scale bar = 10 μm.

**Supplementary Figure 2**

(A) Fluorescent image of testes overexpressing mCherry-CID (CID_OE, red), CAL1-YFP (CAL1_OE, yellow) or CAL1-YFP together with mCherry-CID (CAL1-CID_OE). For CID_OE, nuclei are co-stained with DAPI (cyan). CAL1-YFP_OE and CAL1-CID_OE are co-stained with anti-FIBRILLARIN antibody (cyan) to mark nucleoli. Scale bar = 10 μm. Zoom inset shows a nucleus (white arrow) with the expected localisation pattern for mCherry-CID, CAL1-YFP or CAL1-YFP together with mCherry-CID. Scale bar = 1 μm.
(B) Fluorescent image of mature sperm (stained for DAPI, cyan) in *nanos-GAL4*, CID_OE, CAL1_OE and CAL1-CID_OE lines. Scale bar = 10 μm.

**Supplementary Figure 3**

(A) Immunofluorescent image of testes in *nanos-GAL4*, CID_OE, CAL1_OE and CAL1-CID_OE lines stained for the hub marker ARMADILLO (Arm, red) and DNA is stained with DAPI (cyan). Scale bar = 10 μm.
(B) Quantitation of the number of DAPI-dense nuclei per hub (n=30) in *nanos-GAL4*, CID_OE, CAL1_OE and CAL1-CID_OE lines. ***p<0.001, ns non-significant, error bars = SEM.
(C) Quantitation of the hub diameter in µm (n=30) in *nanos-GAL4*, CID_OE, CAL1_OE and CAL1-CID_OE lines. ****p<0.0001, ns non-significant, error bars = SEM.
(D) Quantitation of total number of GSCs or germ cells up 2CC in the control *nanos-GAL4*, and in CAL1_OE after 10-days. ****p<0.0001, *p<0.05, error bars = SEM.
(E) Ratio of all germ cells up to 2CC divided by the number of GSCs the control *nanos-GAL4*, and in CAL1_OE testes (n=25) after 10-days. *p<0.05, error bars = SEM.
(F) Representative image from *nanos-GAL4* control testis stained with the anti-ɣ-tubulin antibody to mark centrosomes together with a hub marker (red), VASA (grey) and DAPI (cyan). Examples of orientated and misoriented centrosomes in GSCs are indicated. Scale bar = 10 μm.
(G) Percentage of GSCs with misorientated centrosomes in *nanos-GAL4* (n=8/133, 6%), CID_OE (n=12/157, 8%), CAL1_OE (n=4/116, 3%), or CAL1-CID_OE (n=16/168, 10%), lines.

**Supplementary Figure 4**

(A) Immunofluorescent image of control *nanos-GAL4* testis or testis overexpressing HA-CENP-A under *nanos-GAL4* control (HA-CENP-C_OE) stained for HA (yellow), pMAD to mark GSCs (magenta) and DAPI (cyan). White dashed circles indicate pMAD-positive GSCs in contact with the DAPI-dense hub. Scale bar = 10 μm.

(B) Immunofluorescent image of control *nanos-GAL4*, HA-CENP-C_OE, CENP-C RNAi and HA-CENP-C; CENP-C RNAi (HA-CENP-C rescue) testis stained for the spectrosome together with the hub (red) and DAPI (cyan), and CID (red) and CENP-C (green). Scale bar = 10 μm.

(C) Quantitation of the ratio of total CID fluorescent intensity (integrated density) between GSC and GB S-phase pairs (n=25-30) in *nanos-GAL4*, mCherry RNAi and CENP-C RNAi. Values 1.4 (for *nanos-GAL4*), 1.34 (for mCherry RNAi) and 2.04 (for CENP-C RNAi) indicate mean fold differences in intensity. Error bars = SEM.

(D) Ratio of all germ cells up to 2CC divided by the number of GSCs in the control *nanos-GAL4*, mCherry RNAi and CENP-C RNAi lines. **$p<0.01$, ns non-significant, error bars = SEM.

(E) Fluorescent image of mature sperm (stained for DAPI, cyan) in *nanos-GAL4*, HA-CENP-C_OE, CENP-C RNAi and HA-CENP-C; CENP-C RNAi (HA-CENP-C rescue) testis. Scale bar = 10 μm.

Supp Figure 1

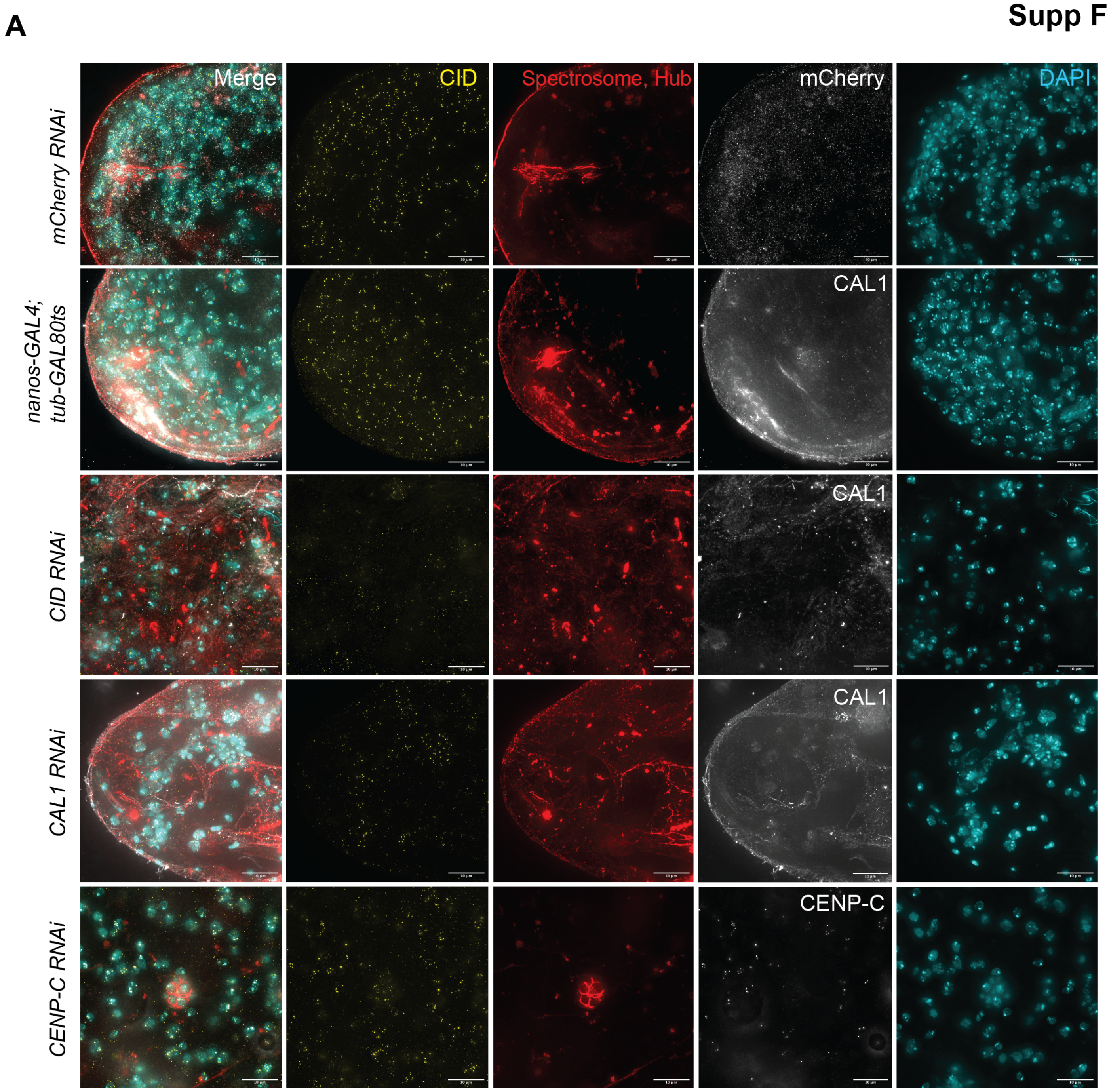


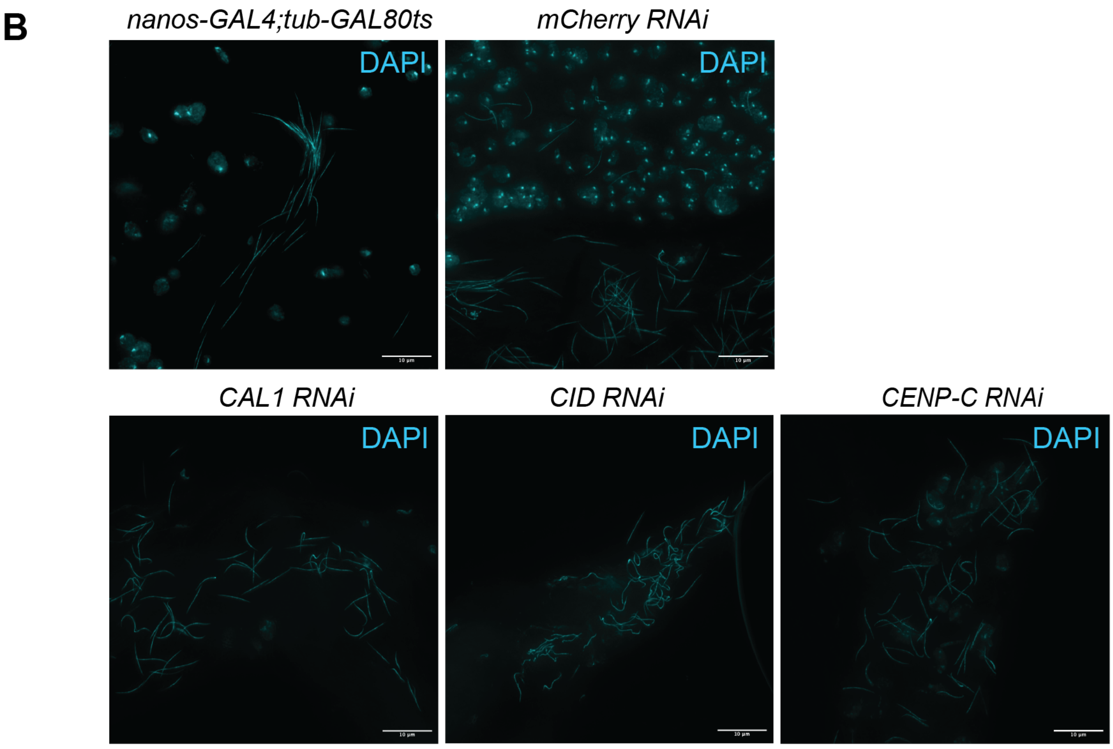

### Supp Figure 2

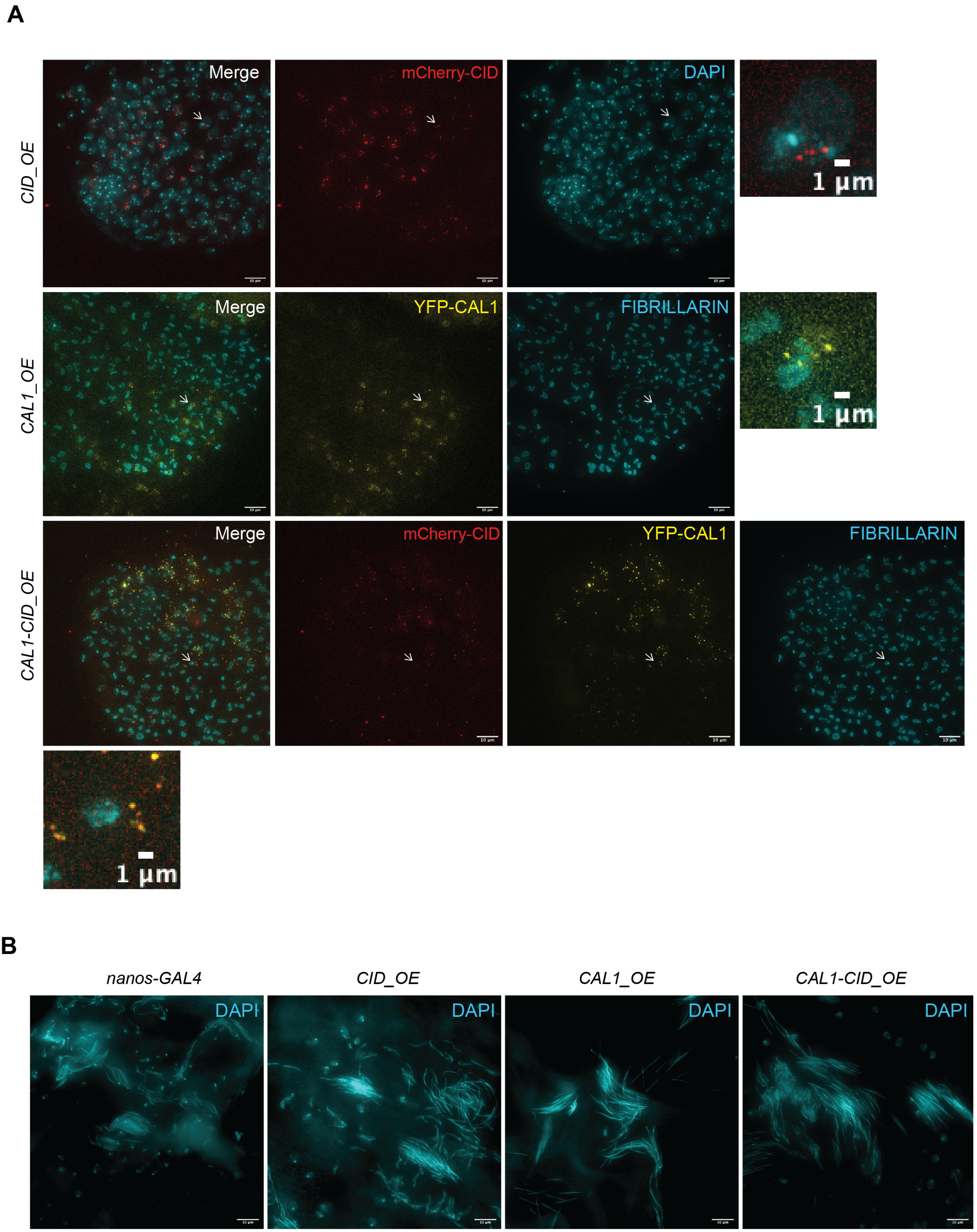

#### Supp Figure 3

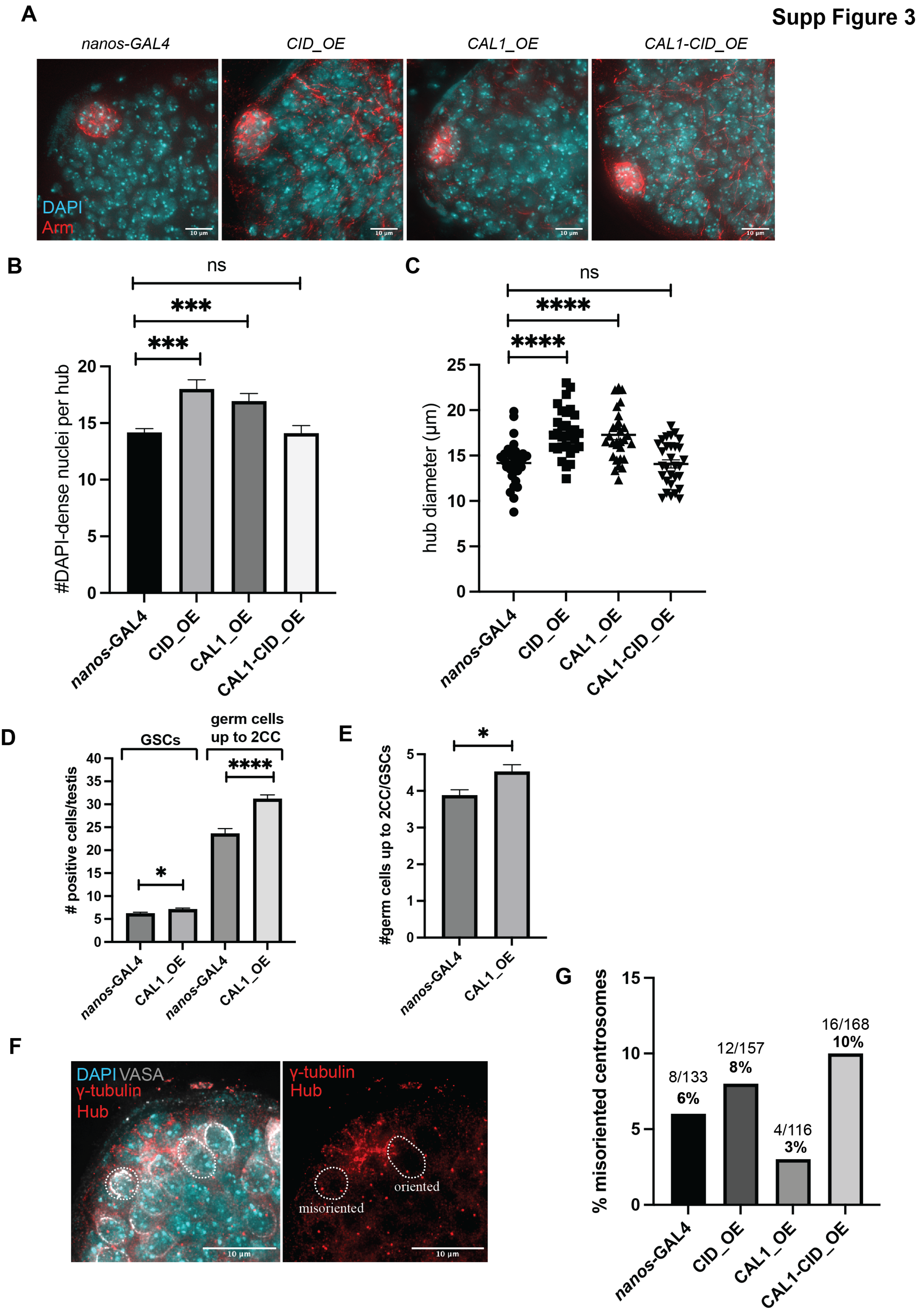

**Supp Figure 4**

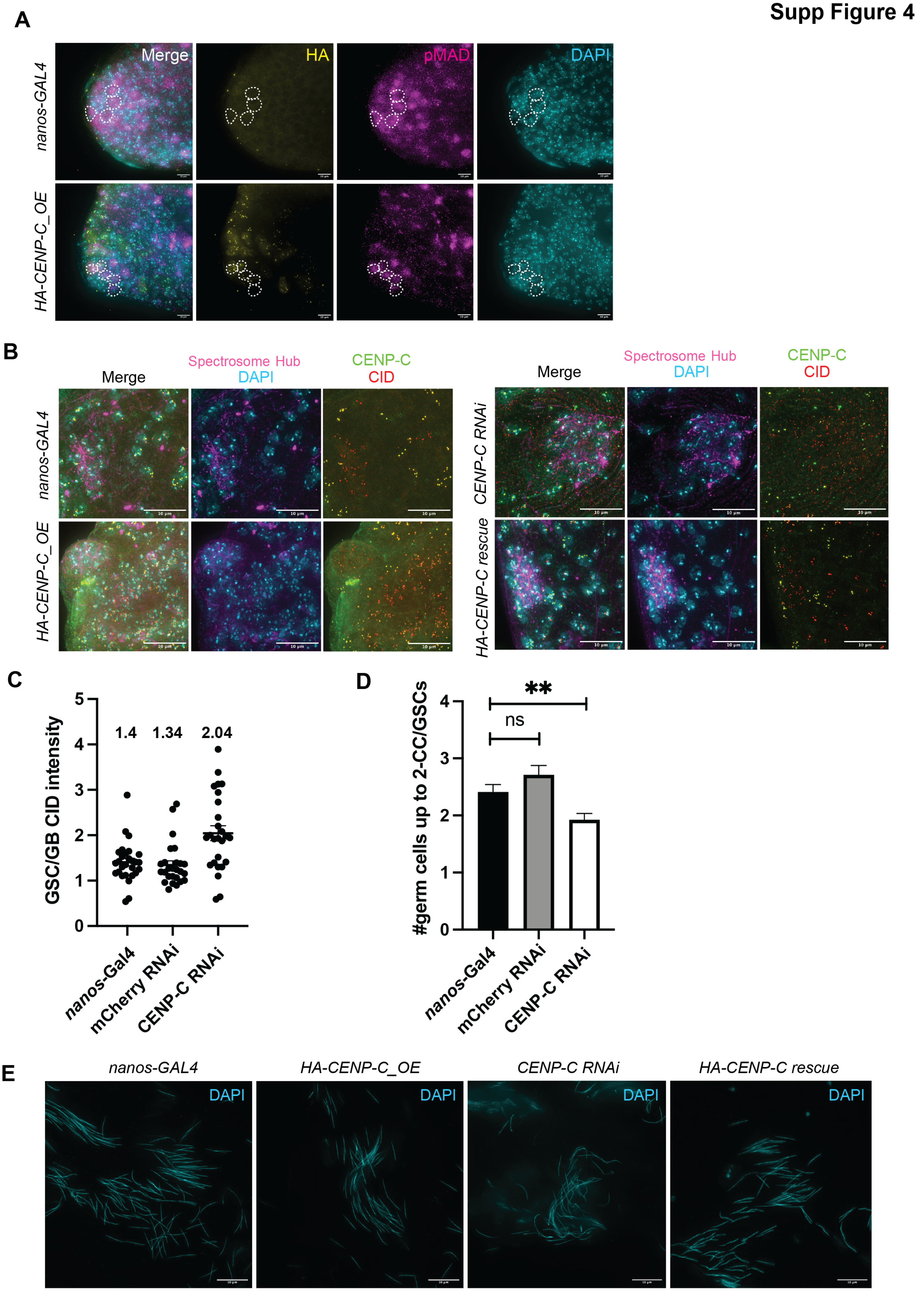
